# Structural basis for polyuridine tract recognition by SARS-CoV-2 Nsp15

**DOI:** 10.1101/2023.11.17.567629

**Authors:** Fumiaki Ito, Hanjing Yang, Z. Hong Zhou, Xiaojiang S. Chen

## Abstract

SARS-CoV-2 non-structural protein 15 (Nsp15) is critical for productive viral replication and evasion of host immunity. The uridine-specific endoribonuclease activity of Nsp15 mediates the cleavage of the polyuridine [poly(U)] tract of the negative-strand coronavirus genome to minimize the formation of dsRNA that activates the host antiviral interferon signaling. However, the molecular basis for the recognition and cleavage of the poly(U) tract by Nsp15 is incompletely understood. Here, we present cryogenic electron microscopy (cryoEM) structures of SARS-CoV-2 Nsp15 bound to viral replication intermediate dsRNA containing poly(U) tract at 2.7-3.3 Å resolution. The structures reveal one copy of dsRNA binds to the sidewall of an Nsp15 homohexamer, spanning three subunits in two distinct binding states. The target uracil is dislodged from the base-pairing of the dsRNA by amino acid residues W332 and M330 of Nsp15, and the dislodged base is entrapped at the endonuclease active site center. Up to 20 A/U base pairs are anchored on the Nsp15 hexamer, which explains the basis for a substantially shortened poly(U) sequence in the negative strand coronavirus genome compared to the long poly(A) tail in its positive strand. Our results provide mechanistic insights into the unique immune evasion strategy employed by coronavirus Nsp15.

## Main

Coronaviruses have evolved a wide array of tactics to evade host antiviral immunity. When host cells detect invading foreign nucleic acids including viral genome and viral replication intermediates, they activate interferon (IFN) signaling via cytoplasmic pattern recognition receptors (PRRs) (1, 2). Remarkable stealth activities exhibited by coronaviruses are facilitated by a series of non-structural proteins (Nsps). For example, coronavirus Nsps possess unique machinery to perform 5′-end capping of their genomic and sub-genomic RNAs to mimic host mRNA, allowing them to escape from the antiviral action of host IFN while hijacking the host ribosome (3). During the initial stage of positive and negative strand RNA genome synthesis, viral RNA-dependent RNA polymerase (RdRp), which consists of Nsp7, Nsp8, and Nsp12, coordinates with viral capping enzymes and their associated cofactors such as Nsp9, Nsp10, Nsp13, Nsp14, and Nsp16. These Nsp proteins form a large replication-transcription complex (RTC) and add the ^7Me^GpppA_2′-OMe_ cap at the 5′-end of the viral RNAs (4-7). This process enables coronaviruses to replicate outside the nucleus without the need to rely on host mRNA capping enzymes.

Another key disguising strategy during the coronavirus replication is carried out by Nsp15, which is conserved among known coronavirus lineages (8, 9). Nsp15 mediates the evasion of cytoplasmic dsRNA detection by host MDA5 and other PRRs in macrophages to suppress the antiviral IFN response (10, 11). Nsp15 is a uridine-specific endoribonuclease and is active on both ssRNA and dsRNA *in vitro* (12-14). Purified Nsp15 cleaves uridine-containing ssRNA with low specificity and dsRNA with high specificity (15, 16). Nsp15 is also a part of the coronavirus RTC assembly and its association with RTC is likely mediated by Nsp8, a known cofactor of RdRp (17-19).

Polyuridine [poly(U)] tract present in the coronavirus negative strand RNA, a transcription product of its positive strand RNA genome, has been identified as one of the physiological targets of Nsp15 (20). The study showed that an endonuclease-deficient mutant of Nsp15 caused the accumulation of dsRNA in the coronavirus-infected hepatocytes and that the negative strand genomic RNA is responsible for this dsRNA formation. They further demonstrated that Nsp15 limits the abundance and length of poly(U) within the negative strand RNA 5′-extension. The transfection of poly(U)-containing synthetic RNA into alpha mouse liver 12 cells caused the activation of MDA5-mediated IFN induction (20). During the coronavirus replication, a short poly(U) sequence in the negative strand is required to initiate the synthesis of a long poly(A) tail (100-130 bp) in the positive strand genome, which in turn is a prerequisite for the negative strand synthesis (21-23). Thus, Nsp15 plays a role in trimming down the initially synthesized poly(U) lead sequence to the optimal length that can suppress dsRNA formation but still serves as a template for poly(A) extension.

X-ray crystallography and cryogenic electron microscopy (cryoEM) studies conducted throughout the SARS-CoV-2 pandemic have advanced our understanding of the substrate binding mechanisms employed by Nsp15 for both ssRNA and dsRNA (24-26). However, the precise molecular mechanisms underlying the recognition of poly(U) by Nsp15 remain unknown. Here we reconstituted a complex of SARS-CoV-2 Nsp15 with a viral replicative dsRNA intermediate containing 3′-end of the viral genome followed by a 20-bp poly(A/U) extension. Cryogenic electron microscopy (cryoEM) and 3D classification revealed Nsp15 hexamer structures at various functional states, including RNA-free and two RNA-bound states. Comparison of these structures show that the poly(U) tract of the sequence is recognized by an Nsp15 hexamer via direct contacts with three subunits in two distinct states. The active site utilizes a base-flipping mechanism to hold the target uracil base in the endonuclease catalytic center for cleavage. Overall, we provide a snapshot of coronavirus Nsp15 bound to the poly(A/U) sequence of its genomic replicative dsRNA intermediate for the evasion of the host antiviral response.

## Results

### CryoEM structures of apo- and RNA-bound Nsp15

To understand the mechanism of poly(U) targeting by SARS-CoV-2 Nsp15, we reconstituted a ribonucleoprotein complex of Nsp15 and a 35-bp dsRNA substrate comprising the final 15-bp of 3′-end of the SARS-CoV-2 genome and 20-bp poly(A/U) extension, which represents a coronavirus genome replication intermediate. Nsp15 C-terminal domain (CTD) belongs to a family of endoribonuclease with two conserved catalytic histidine residues (H234 and H249), which serve as general acid and general base, respectively, to attack the 3′-phosphate of the target uridine (25, 27). To capture the RNA substrate bound to Nsp15 and trap the target uracil base at the nuclease active site, catalytically inactive mutant H234A was used for the reconstitution (Fig. 1A). SARS-CoV-2 Nsp8, a previously hypothesized cofactor of the Nsp15, was also added to facilitate the Nsp15-RNA interaction (17, 18). CryoEM imaging and heterogeneous refinement were performed to evaluate and optimize RNA binding in the reconstituted protein-RNA complex (Fig. S1 and S2). An equimolar ratio of Nsp15:RNA resulted in mostly apo-Nsp15 (Data set 1 in Fig. S2). We found that including an excess amount of RNA with the Nsp15:RNA ratio of 1:10 yielded a complex with the available RNA binding sites within the functional oligomer of Nsp15 occupied by RNA (Data sets 2 and 3 in Fig. S2).

**Fig. 1.**
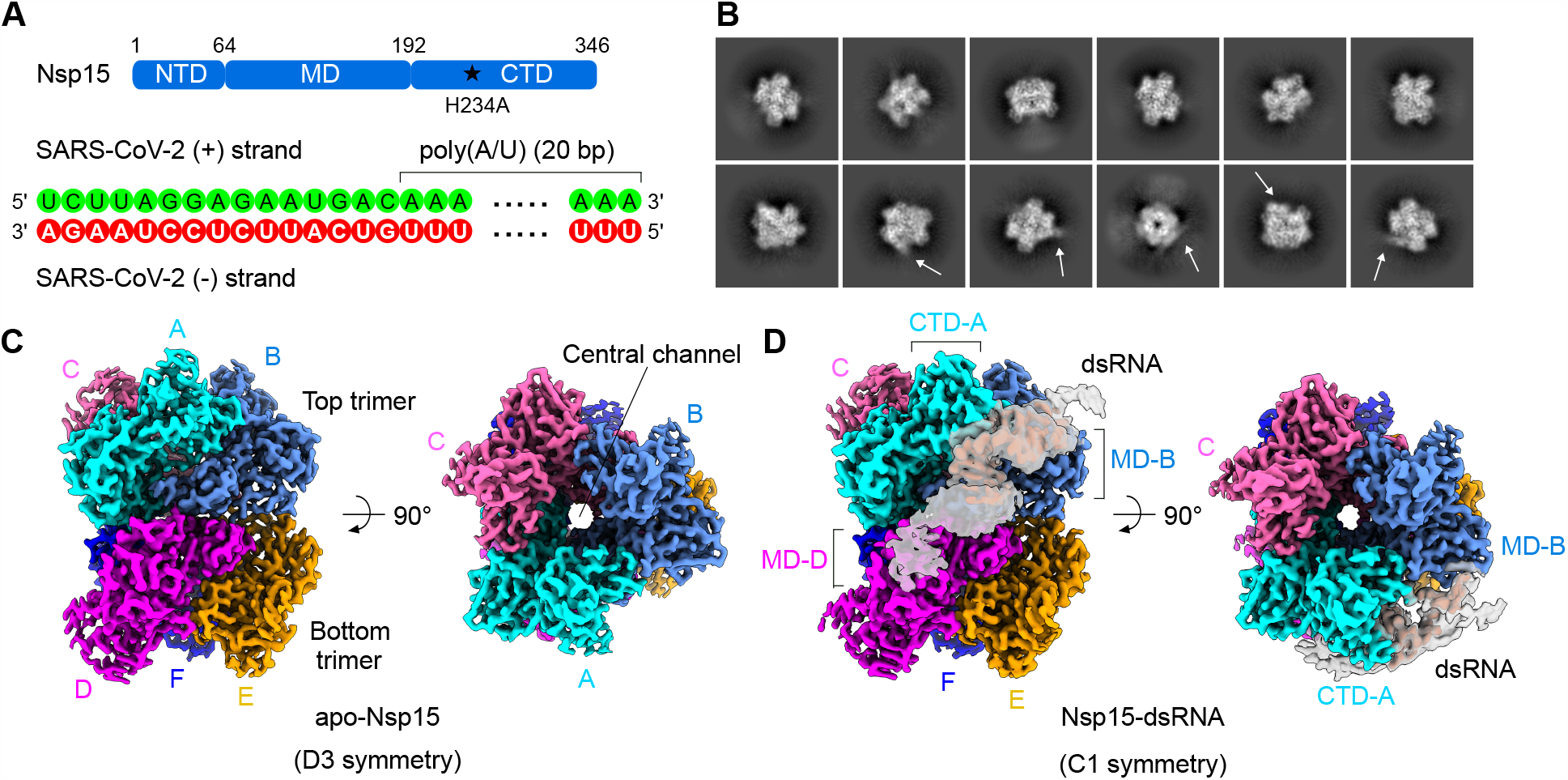
CryoEM reconstructions of apo- and RNA-bound SARS-CoV-2 Nsp15. (A) Schematic representation of the domain organization and construct design of Nsp15 (top). dsRNA substrate sequence used for the reconstitution of Nsp15-RNA complex (bottom). (B) 2D class averages of the reconstituted Nsp15-RNA complex. A subset of the 2D classes shows extra density stemming from the core of the 2D density (indicated by arrows). (C) Two orthogonal views of the cryoEM reconstruction of apo-Nsp15 at 2.3 Å resolution. (D) Two orthogonal views of the cryoEM reconstruction of the Nsp15-RNA complex at 2.7 Å resolution. RNA density is shown in two different isosurface threshold levels (grey and red surfaces) to show both high-resolution and low-resolution features.

2D classification of the reconstituted Nsp15-RNA complex particles in data sets 2 and 3 showed that a subset of 2D class averages has extra helical densities stemming from the core of the 2D densities (Fig. 1B). Multiclass *ab initio* 3D reconstruction showed that 25% of the particles belong to RNA-free apo-Nsp15, and 39% belong to Nsp15 in complex with obvious dsRNA (Fig. S2). The apo-Nsp15 was reconstructed as a homohexamer with D3 symmetry and refined to 2.3 Å resolution (Table S1). The hexamer forms a barrel-like architecture with a central channel, which consists of a head-to-head stack of two trimers (defined as top and bottom)(Figs. 1C, S3, and S4). The RNA-bound form was reconstructed as a homohexamer with clear A-form like RNA duplex density and refined to 2.7 Å resolution (Table S1). dsRNA density is diagonally attached on the outer periphery of the hexameric barrel and occupies a shallow groove between two subunits of the top trimer (defined as subunit A and B)(Figs. 1D, S5, and S6). The nuclease active site centers are located near the subunit interface between the neighboring subunits within the top or bottom trimers and the bound dsRNA fully shields the active site of the subunit A. One end of the dsRNA stretches along a groove between the CTD of subunit A and the middle domain (MD) of subunit B, extending upwards beyond the top trimer. The other end of the dsRNA stretches towards the MD of the subunit D within the bottom trimer. No extra densities were observed in the central channel of the hexamer. Although we added an excess quantity of the substrate RNA (10-fold of Nsp15 in molarity), we only observed Nsp15 hexamer with a single piece of dsRNA as a substrate-bound form. The active sites of the other five subunits are unoccupied. Extensive 3D classification did not yield any 3D classes containing more than one dsRNA piece per Nsp15 hexamer. These observations likely indicate that Nsp15 hexamer is compatible with only one dsRNA substrate at a time.

We noticed that both ends of the bound dsRNA had relatively weak density (Fig. 1D). We therefore hypothesized that there is conformational or compositional variability in the bound RNA structure and that the observed RNA-bound form could be an average of multiple different states. Further heterogeneous 3D refinement revealed that the dsRNA-bound form was subclassified into two states: A class with dsRNA extending towards subunit B in the upper trimer (state 1) and the other class with dsRNA extending towards subunit D in the bottom trimer (state 2). In the state 1 structure, the full 20 A/U pairs are readily visible, including 11-bp preceding the target U at the active site (defined as U_0_) and 9-bp following the U_0_. The state 2 structure also showed the full 20 A/U pairs, including 3-bp preceding the U_0_ and 16-bp following the U_0_ (Figs. 2A and S7). Taken together, these observations indicate that a single piece of dsRNA can engage in the Nsp15 hexamer during poly(U) RNA targeting and that a range of uridines within the poly(U) tract can be targeted for cleavage.

**Fig. 2.**
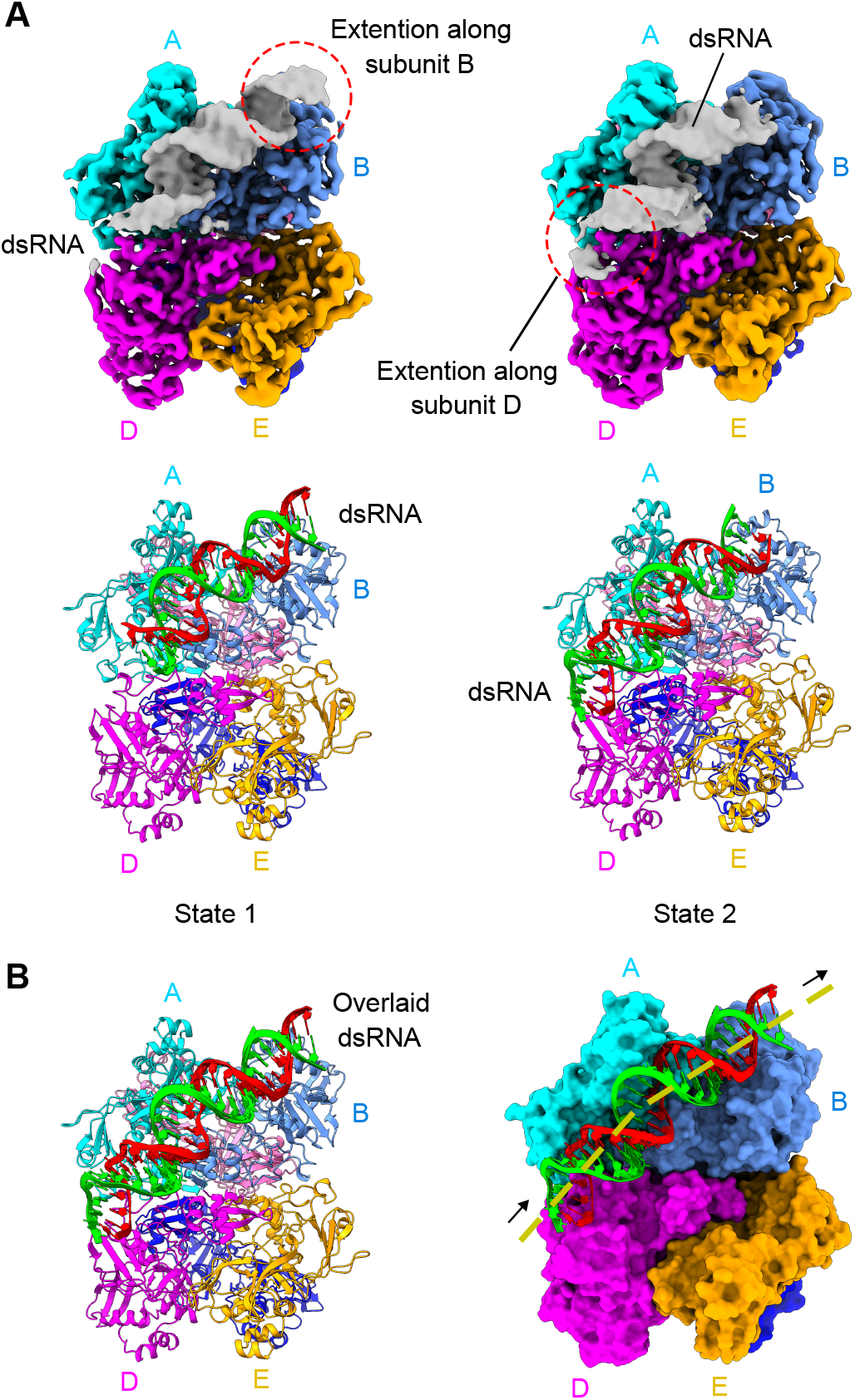
Two distinct states of Nsp15-RNA complex. (A) CryoEM density maps (top) and the corresponding atomic models (bottom) of the Nsp15-RNA state 1 (left) and state 2 (right) structures. The RNA chains in green and red correspond to positive and negative strands of the coronavirus genome, respectively. (B) Overlay of the RNA models from the two states on the consensus Nsp15 structure in the ribbon model (left) and surface model (right). The dotted line indicates the trajectory of the bound dsRNA.

### Poly(U) uracil base flipping at active site center

The dsRNA is firmly held by the Nsp15 hexamer with a slight bent near the groove between the CTD of subunit A and the NTD of subunit B (Fig. 2B). Within the central region of the dsRNA, we observed unpaired bases: one base is flipped outside from the RNA duplex while its complementary counterpart remains orienting inwards within the duplex (Fig. 3A). U/A pair was modeled at this location and the flipped uracil is designated as U_0_. The local resolution of the bound RNA ranges between 2.4 Å to 4.0 Å with the highest resolution around the flipped U_0_ base (Fig. S5). A total of 17 A/U base pairs were confidently built for the consensus Nsp15-RNA structure including the 10-bp preceding the flipped U_0_ and 6-bp following the U_0_. Outside this patch, the base densities are insufficiently featured to distinguish their identity.

**Fig. 3.**
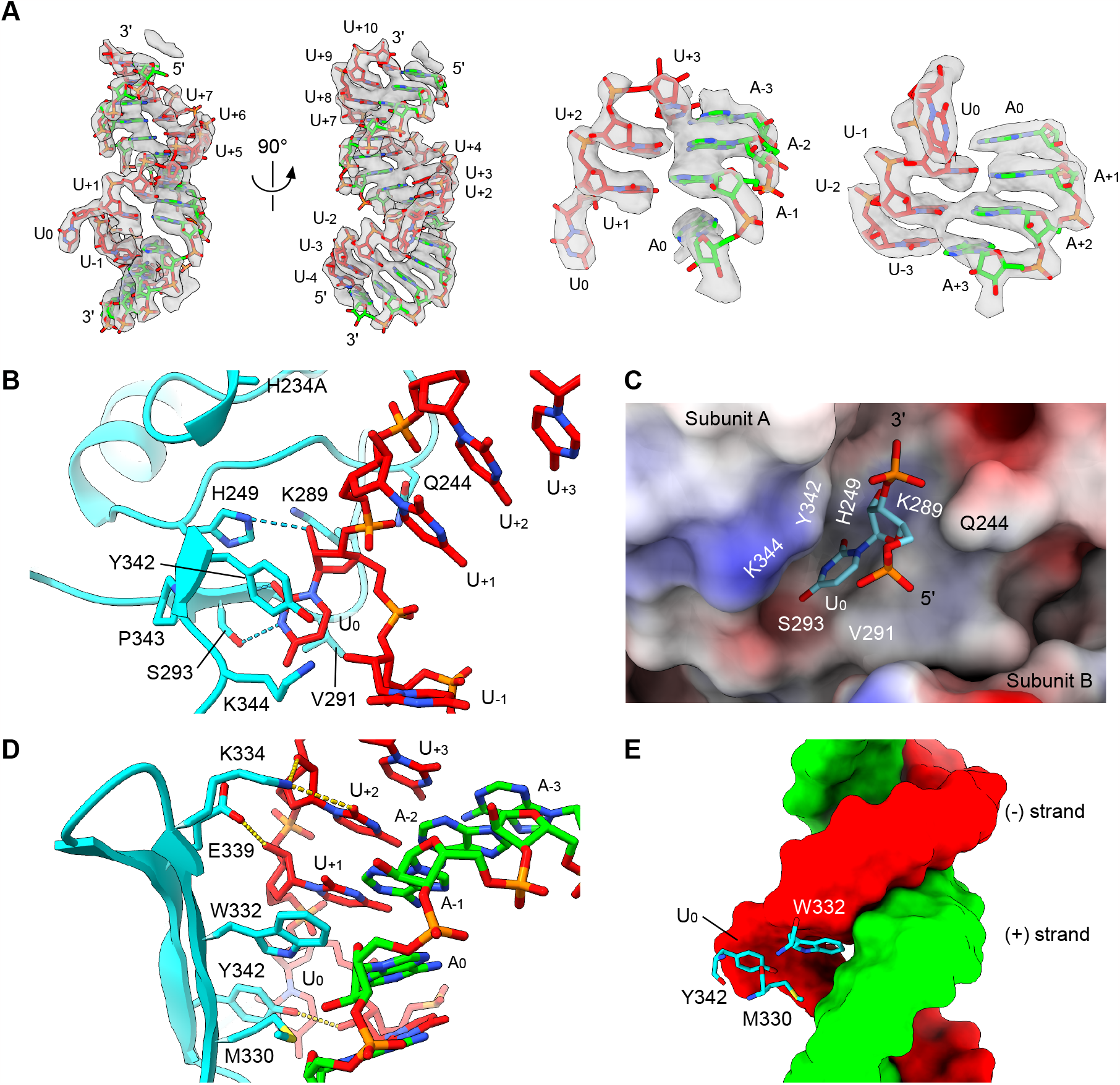
Poly(U) uracil base flipping at nuclease active site center. (A) CryoEM density and the corresponding atomic model of the RNA bound to Nsp15. Two orthogonal views of the bound dsRNA density and its model (left). Density fitting of the RNA model of the U_0_-U_+3_:A_0_-A_+3_ (middle). Density fitting of the RNA model of U_-3_-U_0_:A_+3_-A_0_ (right). (B) Structure of the endonuclease active site center and recognition mechanism of flipped scissile U_0_ base. (C) Surface electrostatic potential of the Nsp15 around the endonuclease active site center and the interactions with the target U_0_ with both 3′- and 5′-phosphates (depicted in sticks). The surface area is colored according to the calculated electrostatic potential from -10.0 kT/e (red) to +10.0 kT/e (blue). (D) Recognition of the open major groove of the dsRNA substrate by Nsp15. W332 is intercalated into the space that would have been occupied by the flipped U_0_ base within the RNA duplex. M330 and Y342 additionally create the hydrophobic surface to facilitate the major groove interaction. Two polar residues K334 and E339 interact with U_+2_ and U_+1_, respectively. (E) The key hydrophobic residues responsible for base-flipping. The side chain of W332 is deeply intercalated into the major groove of the RNA duplex. M330 and Y342 additionally participate in the major groove interaction.

At the endonuclease active site pocket, the pyrimidine ring of the flipped uracil base is sandwiched between two hydrophobic residues Y342 and V291. The aliphatic chain of K344 additionally constitutes this hydrophobic pocket to hold the uracil base. S293 plays a key role in conferring the selectivity for the target uracil as a hydroxyl of the S293 side chain recognizes the N3 atom of the uracil base to form a hydrogen bond. The main chain nitrogen of S293 forms another hydrogen bond with the O2 atom of the uracil base. The catalytic H249 is located near the scissile 3′ -phosphate of U_0_ and its imidazole ring forms a hydrogen bond with the 2′-OH of the ribose ring of U_0_. As expected, the inactivated catalytic histidine (H234A) is located near H249 and the scissile 3′-phosphate. Two polar residues Q244 and K289 additionally surround the 3′-phosphate, thereby stabilizing the position of the target uridine (Fig. 3B and C).

The space that would have been occupied by the U_0_ base within the RNA duplex is partially occupied by W332 from the three-stranded anti-parallel β-sheet of CTD. W332 together with M330 and Y342 from the same β-sheet create a hydrophobic surface and intercalates into the open major groove at this location (Fig. 3D and E). Notably, W332 positions itself directly across the orphan A_0_ base and stabilizes the adjacent U_+1_ base by forming a stacking interaction. These three residues, M330, W332, and Y342, responsible for dislodging the target uracil base are completely conserved across coronaviruses, highlighting their importance (Fig. S8). Two polar residues K334 and E339 near the top edge of the β-sheet interact with the U_+2_ and U_+1_ in the negative strand, respectively. A primary amine of K334 forms hydrogen bonds with the O2 atom of the U_+2_ base, and 2′-OH of the ribose ring of U_+2_. A side chain carboxyl of E339 forms hydrogen bonds with 2′-OH of the ribose ring of U_+1_ (Fig. 3D and E).

### Nsp15-dsRNA interaction at multiple locations

The dsRNA substrate contacts three subunits (subunits A, B, and D) extensively on the sidewall of the Nsp15 hexameric barrel (Fig. 2B). Both the negative and positive strands make substantial contacts with Nsp15 across approximately two and a half turns of the double-stranded helix. Aside from the base pair that involves flipped U_0_, base pairing is well-maintained throughout the duplex. Outside the endonuclease active site pocket anchoring the flipped U_0_ base, subunit A has two additional interface areas on either direction of the active site (Fig. 4A and B). The first interface area includes the first strand of the three-stranded β-sheet in the CTD. A patch of -^314^VSKV^317^-in β13 is in close contact with the orphan nucleotide A_0_ and A_+1_ in the positive strand RNA. S315 forms a hydrogen bond with the 2′-hydroxyl group of the ribose ring of the A_0_, while the backbone nitrogen of V317 forms another hydrogen bond with the 2′-hydroxyl of the ribose ring of A_+1_. The second interface in subunit A is near a patch of -^242^HSQ^244^-within a loop region, which supports the backbone of the positive strand between positions A_-8_ to A_-6_. Subunit B, on the other hand, interacts with the RNA through its MD and NTD (Fig. 4C and D). The surface area comprised of hydrophilic residues S128, E145, S147 and K173 in subunit B’s MD contacts the major groove near U_+8_ to U_+10_ in the negative strand and A_-8_ to A_-6_ in the positive strand (Fig. 4C). Another polar surface comprised of residues K12, D16, Q18, and Q19 in subunit B’s NTD contacts the backbone of the negative strand near U_-4_ to U_-1_ (Fig. 4D). Lastly, Nsp15-RNA state 2 structure has an additional interface in subunit D from the bottom trimer. A hydrophilic surface comprised of the polar residues K110, T112, E113, D132, N136, and R135 in subunit D’s MD contacts the minor groove area near U_-14_, and U_-13_ in the negative strand and A_+15_, and A_+16_ in the positive strand, further stabilizing the bound dsRNA (Fig. 4E).

**Fig. 4.**
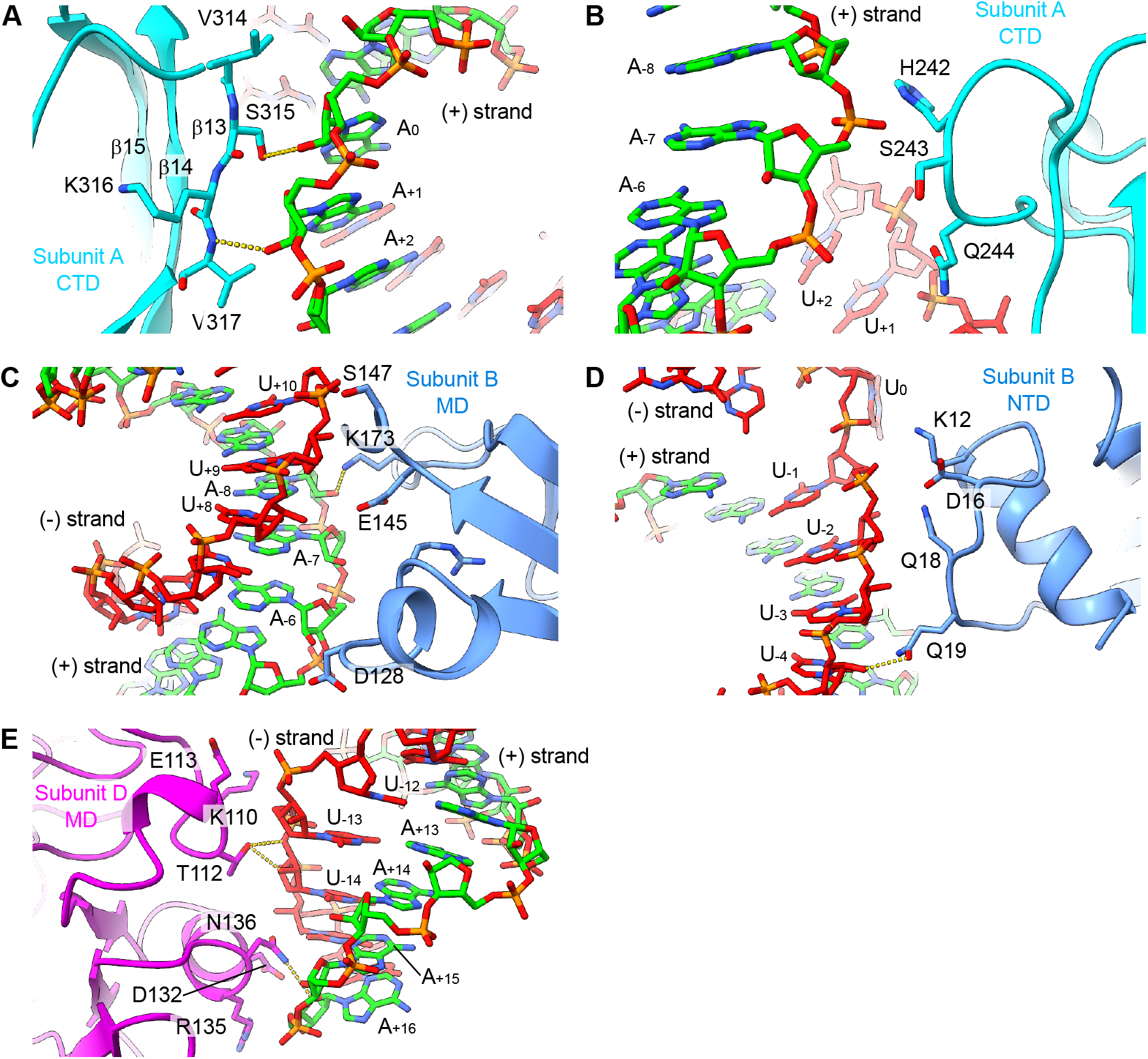
Bound RNA forms extensive interactions with Nsp15 hexamer across three subunits. (A) Interface between a patch of -^314^VSKV^317^-of subunit A’s CTD and A_0_ and A_+1_ of the positive strand. (B) Interface between a patch of -^242^HSQ^244^-of subunit A’s CTD and A_-8_ and A_-7_ of the positive strand and U_+1_ of the negative strand. (C) Interface between subunit B’s MD and U_+10_ of the negative strand and A_-8_, A_-6_ of the positive strand. (D) Interface between subunit B’s NTD and the U_-4_-U_-1_ of the negative strand. (E) Interface between subunit D’s MD and U_-14_-U_-12_ of the negative strand and A_+16_ of the positive strand.

### Structural remodeling by dsRNA binding

Comparison of subunit A structures in its RNA-bound form and its apo-form showed an r.m.s.d. of 0.388 Å, indicating that the binding of RNA does not induce significant global conformational changes to Nsp15 protomer. Yet, the high-resolution structures of both apo- and RNA-bound forms of Nsp15 allowed us to identify notable local structural changes. First, upon binding of RNA, there is a subtle linear shift of β-strands 14 and 15, accompanied by their connecting β-turn (-^334^KDGH^337^-), towards the bound dsRNA. This shift is likely caused by the presence of W332 on the β-strand 14 (Fig. S9A). Second, the C-terminal tail of the subunit A including the terminal residues -^344^KLQ^346^ underwent a structural remodeling upon RNA binding. Superimposition of the apo-Nsp15 and RNA-bound subunit A structures showed that the C-terminal end glutamine residue clashes with the flipped uracil base. In the RNA-bound structure, the ^344^KLQ^346^ patch swung away from the active site pocket, allowing the substrate uracil base to fit into the pocket (Fig. S9B). The rest of the subunits (B to F) did not display any noticeable structural changes upon RNA binding. Interestingly, the cryoEM density for residues W332 and M330 in subunit A, which play key roles in the base-flipping of the target uracil, are more clearly defined than the same residues in the other subunits, indicating that the side chains of these residues are stabilized by the RNA binding (Fig. S9C).

## Discussion

Nsp15 is one of the indispensable components of the non-structural proteins that are initially expressed as a large polyprotein and subsequently proteolytically cleaved to yield individual functional enzymes. Nsp15 plays a pivotal role in facilitating viral infection. Inactivation of its endonuclease activity severely compromises the virus fitness, thereby making it a promising therapeutic target (10, 11). Our understanding of Nsp15’s physiological role remains limited due to a lack of information about its array of physiological substrates, the regulatory mechanisms governing its endonuclease activity, and its association with other viral or host proteins. Nsp15 targets the extended poly(U) lead sequence at the 5′-end of the coronavirus negative strand genome. This activity curtails the accumulation of dsRNA, effectively suppressing the activation of the antiviral response mediated by the cytoplasmic dsRNA sensor MDA5 (20). Our cryoEM structures of SARS-CoV-2 Nsp15 in complex with its substrate RNA provide direct evidence of its association with poly(U) in its double-stranded form, which represents a replication intermediate during the initial synthesis of poly(U) tract using positive-strand poly(A) tail as a template. Our results also show that the Nsp15-RNA complex exists at least in two distinct states, indicating that a range of uridines within the poly(U) tract can be recognized for cleavage in the reconstituted system. The target uracil base is flipped out from the duplex and is captured at the endonuclease active site pocket. The hydrophobic residues W332 and M330 are intercalated into the open major groove of the RNA duplex and are likely responsible for dislodging the target uracil base.

Prior crystallographic and cryoEM studies of Nsp15 have provided insights into its binding to both ssRNA and dsRNA (16, 24, 26). Overall dsRNA binding mode observed in our Nsp15-RNA structures resembles that observed in the recently reported structure of SARS-CoV-2 Nsp15 bound to a 52-bp dsRNA (26). A synthetic dsRNA duplex adopted from a substrate of the *Drosophila* Dicer-2 was used in earlier studies (26, 28), in contrast to the poly(A/U)-containing physiological dsRNA substrate used in the current study. The observed target uracil base recognition mode in our structure is consistent with both short ssRNA-bound and dsRNA-bound structures (24, 26). Notably, the structure with 52-bp dsRNA shows the adenine at the +1 position of the target strand. The structure presented in the current study shows that uridine can also be readily accommodated at this location. Interestingly, the comparison of the structures around the target U_0_ base shows that the position of W332 is tuned to promote the interactions by stacking the indole ring with the U_+1_ base while the flipped U_0_ base remains precisely at the same position.

Nsp15 is part of the large coronavirus RTC, which is responsible for RNA-dependent RNA transcription and 5′-capping (17, 29). The RTC is formed by a series of Nsps including Nsp7, Nsp8, Nsp9, Nsp10, Nsp12, Nsp13, Nsp14, and Nsp16 (5, 17, 19). The association of Nsp15 with RTC provides a timely advantage to viral infection, as Nsp15 can trim the poly(U) tract immediately after its synthesis by RdRp thus minimizing the duration of the presence of unprocessed long poly(U) in the infected cells. Within the RTC, Nsp8 serves as a cofactor of Nsp15, which may jointly hold the substrate dsRNA (30). Despite our attempt to assemble a complex of Nsp15 with Nsp8 in the presence and absence of the substrate RNA, we were unable to observe complex formation between them under the conditions tested. Further study is needed to understand the potential role of Nsp15 in the RTC and their native complex formation during coronaviral replication.

In summary, our structures of the Nsp15-RNA complex in two states reveal direct interactions between Nsp15 and poly(A/U) RNA, its only known physiological substrate. These structures offer snapshots that inform how SARS-CoV-2 camouflages itself in infected cells to escape the host detection of viral RNA. Given the high sequence conservation of endoribonuclease among known coronavirus lineages and SARS-CoV-2 variants known to date, targeting Nsp15 activity may be a promising therapeutic strategy against the current and future SARS-CoV-2 variants. Inhibitors of the Nsp15 activity would preserve the natural innate immune responses against dsRNA derived from the coronavirus genome, giving rise to broad-spectrum anti-viral drugs.

## Methods

### Plasmids

Nsp15 from SARS-CoV-2 isolate WA-CDC-02982586-001/2020 (GenBank: MN985325.1, residues 1-346) with His_6_-tag at N-terminus was cloned into Champion™ pET SUMO vector by excluding SUMO fusion tag, and Nsp8 from the same SARS-CoV-2 isolate (residues 1-198) was cloned into pET28a vector with His_6_-tag at N-terminus. Cloning and mutagenesis were performed with In-Fusion cloning and PrimeSTAR mutagenesis (Clontech) by following the manufacturer’s instructions. The sequences of all the constructs were verified by Sanger DNA sequencing (Azenta Life Sciences). The multiple sequence alignments were generated with Linnaeo (https://github.com/beowulfey/linnaeo).

### Protein expression and purification

His_6_-Nsp15 catalytically inactive mutant H234A and wild-type His_6_-Nsp8 expression vectors were transformed into the *E. coli* strains BL21(DE3). The *E. coli* cells harboring the expression vectors were grown in LB medium at 37°C until the OD_600_ reaches 0.6. The recombinant proteins were induced by 0.2 mM isopropyl β-D-1-thiogalactopyranoside (IPTG) at 16°C for 18 hours.

For Nsp15, the cell pellets were resuspended with the buffer (25 mM HEPES-NaOH (pH 7.5), 500 mM NaCl, and 0.5 mM TCEP) containing RNase A (0.1 mg/ml, Qiagen), lysed by sonication, and cellular debris was removed by centrifugation. The supernatant containing the His_6_-Nsp15 was loaded onto the Ni-NTA agarose column (Qiagen). The nickel column was extensively washed with wash buffer (25 mM HEPES-NaOH (pH 7.5), 500 mM NaCl, 50 mM imidazole, and 0.5 mM TCEP) and the protein was eluted with elution buffer (25 mM HEPES-NaOH (pH 7.5), 500 mM NaCl, 500 mM imidazole, and 0.5 mM TCEP). The eluted proteins were concentrated and subjected to Superdex 200 Increase 10/300 GL column (Cytiva) equilibrated with the buffer (25 mM HEPES-NaOH (pH 7.5), 150 mM NaCl, and 0.5 mM TCEP). The peak fractions corresponding to the hexamer form were collected and concentrated for cryoEM study.

For Nsp8, the cell pellets were resuspended with the buffer (20 mM Tris-HCl (pH 8.0), 500 mM NaCl, and 0.5 mM TCEP) containing RNase A (0.1 mg/ml, Qiagen), lysed by sonication, and cellular debris was removed by centrifugation. The supernatant containing the His_6_-Nsp8 was loaded onto the Ni-NTA agarose column (Qiagen). The nickel column was extensively washed with wash buffer (20 mM Tris-HCl (pH 8.0), 500 mM NaCl, 20 mM imidazole, and 0.5 mM TCEP) and the protein was eluted with elution buffer (20 mM Tris-HCl (pH 8.0), 500 mM NaCl, 300 mM imidazole, and 0.5 mM TCEP). The eluted proteins were concentrated and subjected to Superdex 200 Increase 10/300 GL column (Cytiva) equilibrated with the buffer (20 mM Tris-HCl (pH 8.0), 250 mM NaCl, and 0.5 mM TCEP). The peak fractions were collected and concentrated for cryoEM study. Protein purity was assessed by SDS-PAGE at each purification step.

### Negative-stain EM

5 μl of 0.02 mg/ml purified Nsp15 sample was applied onto glow-discharged ultrathin formvar/carbon supported copper 400-mesh grids (Electron Microscopy Sciences), blotted and stained with 2.0% uranyl acetate. Negative-stained grids were imaged on a Talos F200C transmission electron microscope (Thermo Fisher Scientific) operated at 200 kV.

### CryoEM data acquisition

Three data sets were collected in separate TEM sessions. For reconstitution of Nsp15-RNA complex, pre-annealed dsRNA (chain 1: 5′-rUrCrUrUrArGrGrArGrArArUrGrArCrArArArArArArArArArArArArArArArArArArArA-3′, chain 2: 5′-rUrUrUrUrUrUrUrUrUrUrUrUrUrUrUrUrUrUrUrUrGrUrCrArUrUrCrUrCrCrUrArArGrA-3′) substrate was synthesized (Integrated DNA Technologies). For data set 1, Nsp15, Nsp8, and dsRNA were mixed by 1:0.5:1 molar ratio (7.5 μM Nsp15, 3.75 μM Nsp8, and 7.5 μM dsRNA) in a buffer (25 mM HEPES-NaOH, 150 mM NaCl, pH 7.5). For data sets 2 and 3, Nsp15, Nsp8, and dsRNA were mixed by 1:1:10 molar ratio (7.5 μM Nsp15, 7.5 μM Nsp8, and 75 μM dsRNA) in a buffer with lower salt (25 mM HEPES-NaOH, 100 mM NaCl, pH 7.5). The mixture was incubated on ice for 30-60 min before plunge-freezing. 4 ul aliquots of the mixture were applied to UltrAu foil R1.2/1.3 gold 300-mesh grids (Electron Microscopy Sciences). Grids were then blotted and vitrified in liquid ethane using Vitrobot Mark IV (Thermo Fisher Scientific). CryoEM data was collected in a Glacios (Thermo Fisher Scientific) equipped with Falcon-4 direct electron detector operated at 200 kV in electron counting mode. Movies were collected at a nominal magnification of 150,000× and a pixel size of 0.92 Å in EER format. A total dose of 52 e^-^/Å^2^ per movie was used with a dose rate of 5-6 e^-^/Å^2^/sec. 7,268, 10,001, and 5,150 movies were recorded for the data set 1, 2, and 3, respectively, by automated data acquisition with EPU.

### CryoEM data processing

The movies from three data sets were imported into cryoSPARC software package (31) and subjected to patch motion correction and CTF estimation in cryoSPARC. For data set 1, reference-free manual particle picking in a small subset of data was performed to generate 2D templates for auto-picking. A total of 2,478,629 particles were picked initially, extracted, and down-sampled by a factor of 4, on which 2D classification was performed. 1,961,839 particles from 2D class averages were selected and re-extracted with full-resolution. 3D *ab initio* reconstruction was then performed to generate three initial volumes. A single dominant class containing 68% of the particles showed a feature of hexamer form Nsp15. Further classification did not yield any 3D classes containing RNA or Nsp8 density. For data sets 2 and 3, 2,983,555 and 1,730,135 particles were picked initially by using the templates generated from the data set 1, extracted, and down-sampled by a factor of 4, on which 2D classification was performed. Additional RNA densities were present in a subset of 2D classes, which are not present in the data set 1. 1,330,310 particles (data set 2) and 1,119,650 particles (data set 3) from 2D class averages were selected and re-extracted with full-resolution. 3D *ab initio* reconstruction was then performed to generate three initial volumes. A class containing 39% of the particles in each data set showed a clear feature of dsRNA density attached to the Nsp15 hexameric barrel. A class containing 37-38% of the particles in each data set showed a feature of hexamer form, which is similar to the class observed in the data set 1. The 3D classes representing hexamer form with no obvious RNA densities from the three data sets were combined and non-uniform refinement (32) was performed with D3 symmetry to yield the final 2.3 Å resolution map. The 3D classes representing RNA-bound form from the data sets 2 and 3 were combined and non-uniform refinement was performed with C1 symmetry to yield the final 2.7 Å resolution map. We noticed that RNA density in the 3D map was anisotropic, and the map could be an average of different conformational states. To further classify into possible different classes, heterogeneous refinement was performed to yield four classes. A class containing 24% of the particles showed strong RNA density along the top trimer (state 1) while two classes containing 53% of the particles showed strong RNA density along the bottom trimer (state 2). Each state was subjected to non-uniform refinement to yield the final 3.3 Å (state 1) and 3.1 Å (state 2) resolution maps, respectively. Any additional classification did not yield 3D classes with Nsp8, or Nsp15-RNA complex with more than one dsRNA bound to the Nsp15 hexamer. All resolution evaluation was performed based on the gold-standard criterion of Fourier shell correction (FSC) coefficient at 0.143 (33).

### Model building and refinement

An atomic model derived from crystal structure of SARS-CoV-2 Nsp15 (PDB ID: 6VWW)(27) was docked into the cryoEM map of apo-Nsp15 using UCSF Chimera (34). The apo-Nsp15 model was refined with the phenix.real_space_refine module in Phenix, with secondary structure restraints and geometry restraints (35, 36). The atomic models went through iterative cycles of manual adjustment in COOT (37) and real-space refinement in Phenix (38). For Nsp15-RNA complex consensus form, standard A-form double-stranded RNA was generated and docked together with apo-Nsp15 model into the cryoEM map using UCSF Chimera. The RNA model was manually adjusted while keeping proper RNA geometry using COOT. Nsp15-RNA complex states 1 and 2 models were built based on the consensus form by extending and refining the RNA strands. The final atomic models were validated using the comprehensive cryoEM validation tool implemented in Phenix (Table S1) (39). All structural figures were generated with UCSF ChimeraX (40).

## Acknowledgments

Electron microscopy data were collected at the Core Center of Excellence in Nano Imaging (CNI) at USC. CryoEM data was computed at Center for Advanced Research Computing (CARC) at USC. We thank Htet Khant, Carolyn Marks, and John Curulli for assisting with the operation and maintenance of transmission electron microscopes at CNI, Tomek Osinski for assisting with computing work at CARC, and Cornelius Gati for advice on cryo-EM sample preparation and data processing. This work was funded by NIH grant R01AI150524 to X.S.C. and R01GM071940 to Z.H.Z..

## Author contributions

Z.H.Z. and X.S.C. supervised the project and acquired the funding. F.I. conceived the project and designed the experiments. F.I. and H.Y. generated clones and purified the proteins. F.I. performed the cryoEM grid screening, data collection, image processing, and atomic model building. F.I. wrote the manuscript with inputs from all authors.

