## Supplemental information for "Structural basis for polyuridine tract recognition by SARS-CoV-2 Nsp15"

Table S1. CryoEM data collection, refinement, and validation statistics

| | Apo-Nsp15 | Nsp15-RNA consensus | Nsp15-RNA state 1 | Nsp15-RRNA state 2 |
|---|---|---|---|---|
| **Data collection** | | | | |
| Magnification | 150,000 | 150,000 | 150,000 | 150,000 |
| Voltage (kV) | 200 | 200 | 200 | 200 |
| Electron exposure ($e^-/Å^2$) | 52 | 52 | 52 | 52 |
| Defocus range (μm) | -0.8 to -3.0 | -0.8 to -3.0 | -0.8 to -3.0 | -0.8 to -3.0 |
| Pixel size (Å) | 0.92 | 0.92 | 0.92 | 0.92 |
| Symmetry imposed | D3 | C1 | C1 | C1 |
| Initial particle images | 7,192,319 | 4,713,690 | 4,713,690 | 4,713,690 |
| Final particle images | 2,294,976 | 961,569 | 227,075 | 511,627 |
| Map resolution (Å) | 2.33 | 2.67 | 3.25 | 3.13 |
| FSC threshold | 0.143 | 0.143 | 0.143 | 0.143 |
| Map resolution range (Å) | 2.0 - 2.5 | 2.2 - 4.0 | 2.4 - 5.0 | 2.3 - 5.0 |
| **Refinement** | | | | |
| Initial model used | PDB 6VWW | Apo-Nsp15 | Nsp15-RNA consensus | Nsp15-RNA consensus |
| Model resolution (Å) | 2.4 | 2.8 | 3.4 | 3.2 |
| FSC threshold | 0.5 | 0.5 | 0.5 | 0.5 |
| Map sharpening B factor ($Å^2$) | 112.1 | 104.3 | 99.5 | 117.9 |
| No. non-hydrogen atoms | 16530 | 17244 | 17498 | 17664 |
| Protein residues | 2076 | 2076 | 2076 | 2076 |
| Nucleotides | 0 | 34 | 46 | 54 |
| *B*-factors | | | | |
| Protein | 41.65 | 47.82 | 60.80 | 40.61 |
| Nucleotide | - | 7.45 | 80.60 | 32.35 |
| R.m.s. deviations | | | | |
| Bond lengths (Å) | 0.004 | 0.003 | 0.003 | 0.002 |
| Bond angles (°) | 0.953 | 0.523 | 0.562 | 0.490 |
| Validation | | | | |
| MolProbity score | 1.37 | 1.41 | 1.42 | 1.21 |
| Clash score | 6.23 | 7.53 | 7.63 | 7.89 |
| Poor rotamers (%) | 1.08 | 0.65 | 0.43 | 0.27 |
| Ramachandran plot | | | | |
| Favored (%) | 98.50 | 98.59 | 98.45 | 98.79 |
| Allowed (%) | 1.50 | 1.41 | 1.55 | 1.21 |
| Disallowed (%) | 0.00 | 0.00 | 0.00 | 0.00 |

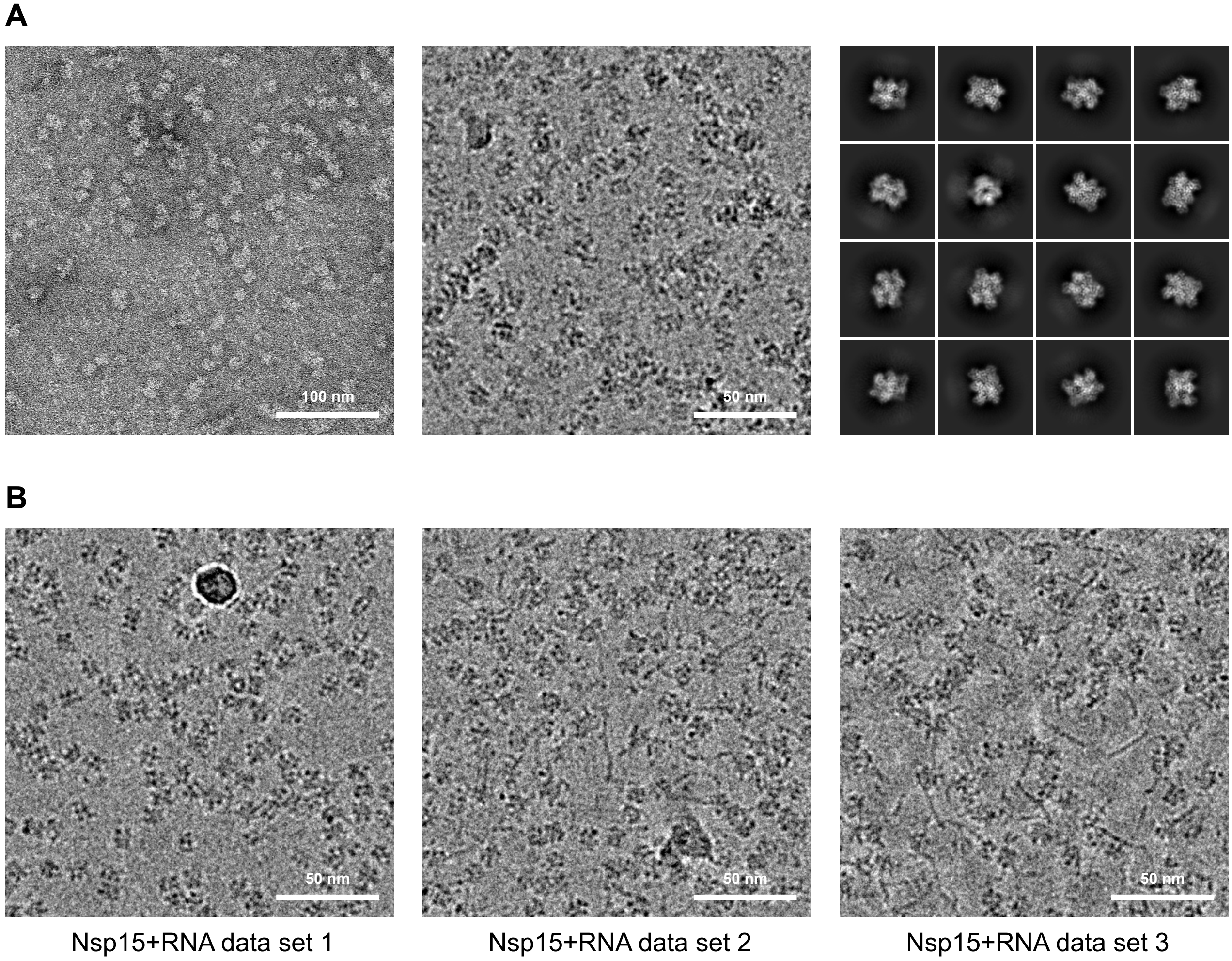


**Fig. S1.** Raw electron microscopy images of Nsp15. (A) Representative negative stain EM raw image (left), cryoEM raw image (middle), and cryoEM 2D class averages (right) of the apo-Nsp15. (B) Representative cryoEM raw images of Nsp15+RNA data sets.

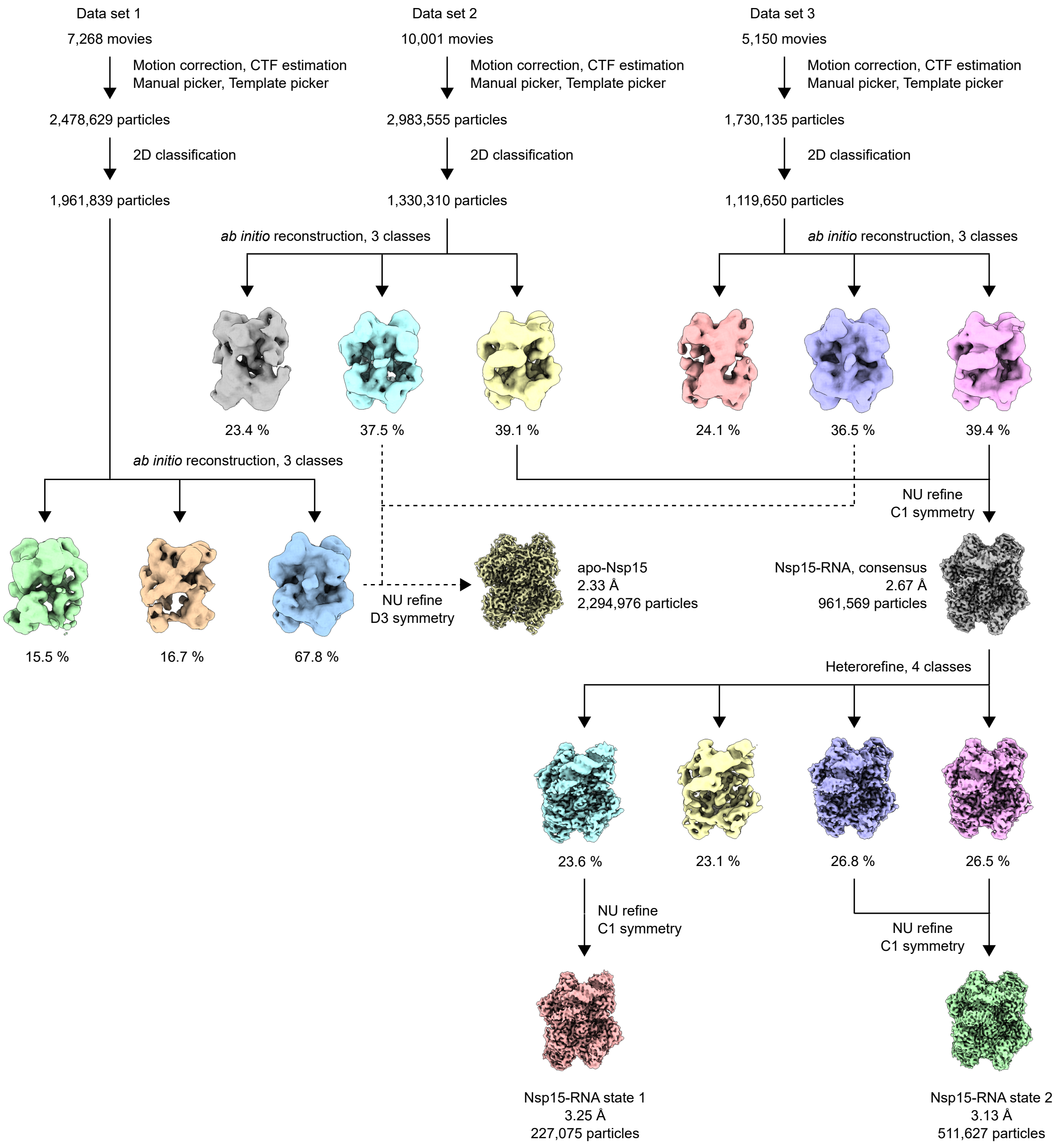


**Fig. S2.** Workflow and intermediate results of cryoEM image processing and 3D reconstruction of the apo-Nsp15 and the Nsp15-RNA complex.

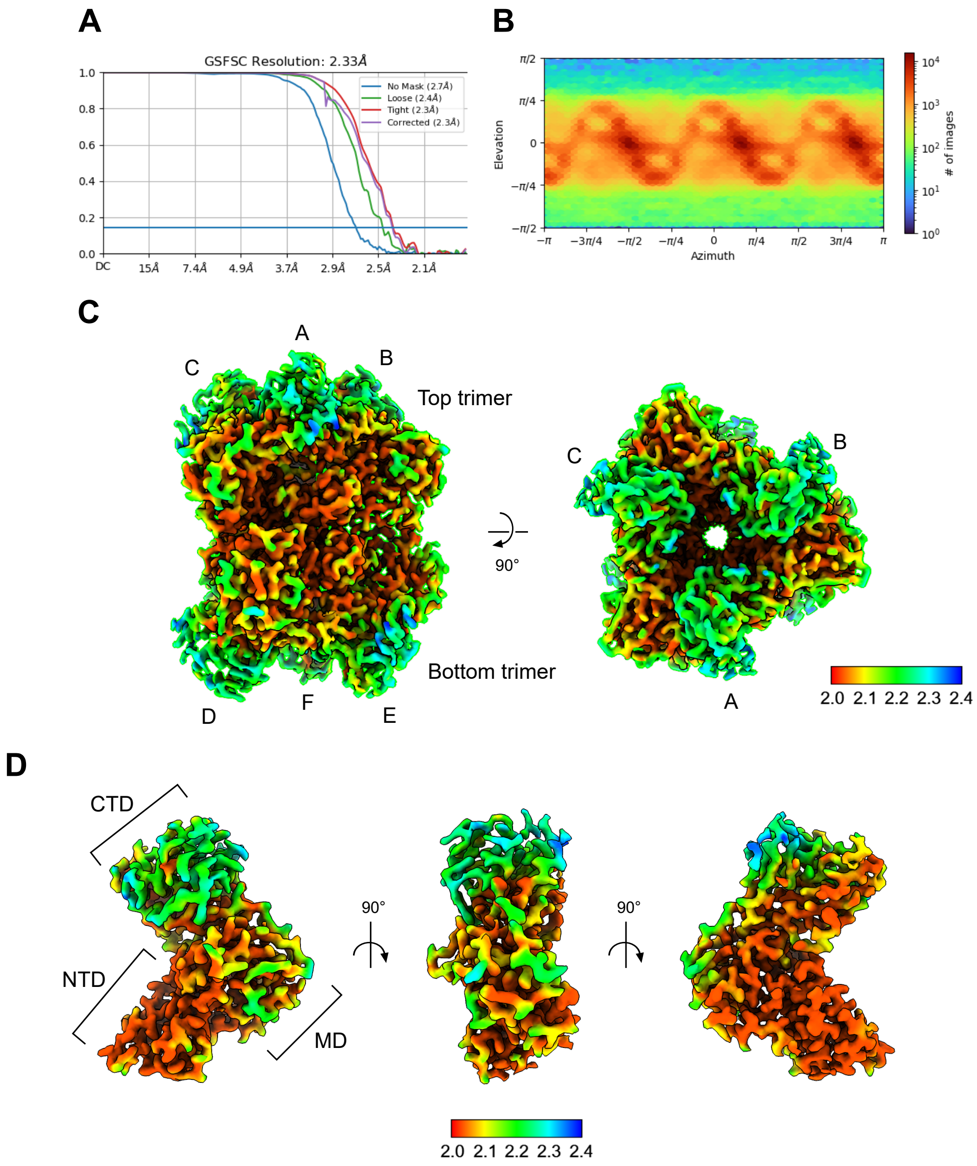


**Fig. S3.** Evaluation of the 3D reconstruction of apo-Nsp15. (A) Global resolution estimation of apo-Nsp15. (B) Angular distribution plot of the particles of apo-Nsp15. (C and D) Local resolution evaluation of the apo-Nsp15 hexamer (C) and one of its protomers (D). Resolution estimation is based on the gold standard Fourier shell correlation (FSC) coefficient of 0.143 criteria.

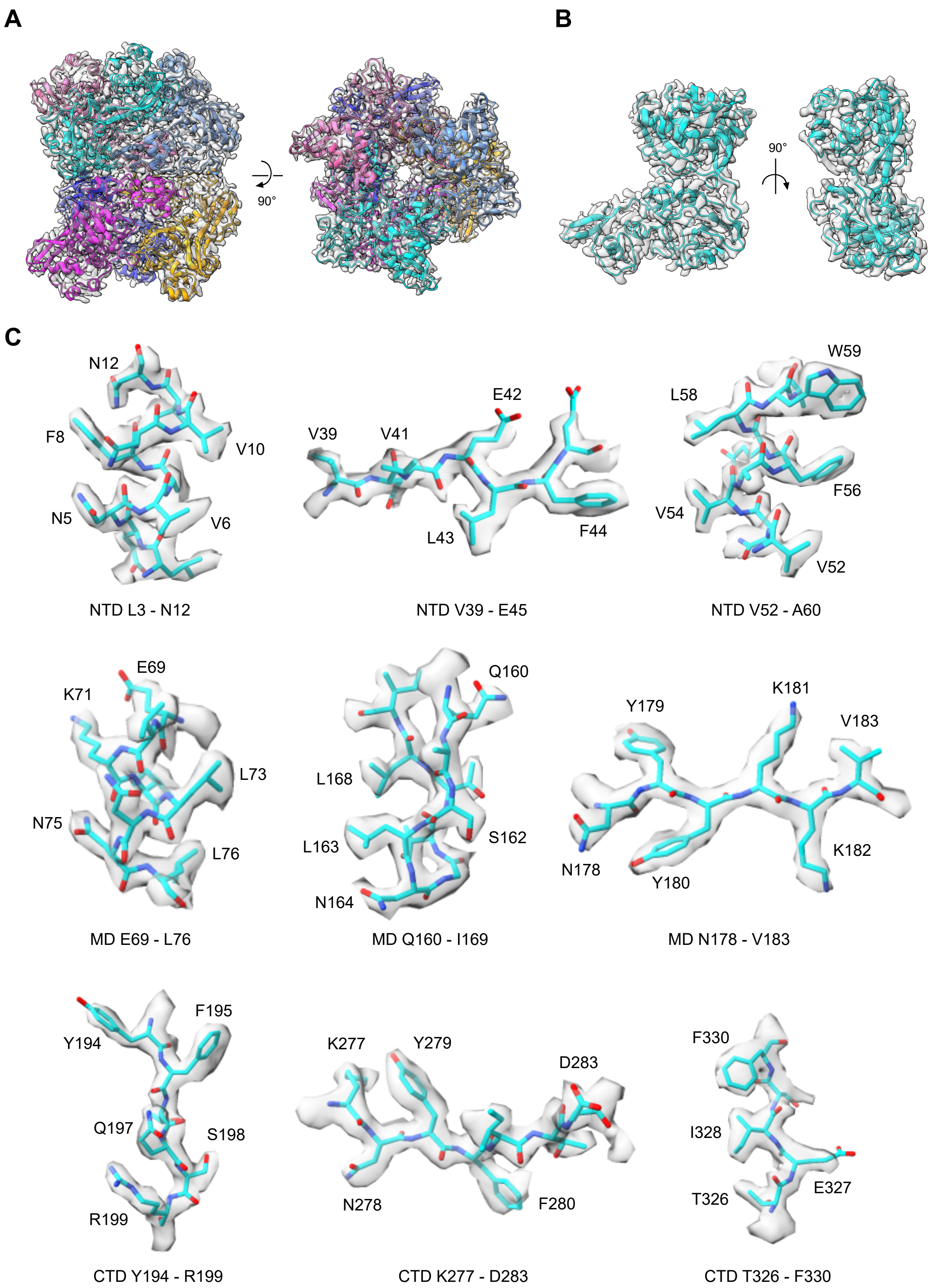


**Fig. S4.** Regions of the cryoEM density maps superposed with atomic models of the components of apo-Nsp15. Overall cryoEM density of apo-Nap15 hexamer (A), segmented cryoEM densities of representative protomer of apo-Nsp15 (B), and its local regions (C) are shown as semi-transparent surfaces superposed with atomic models of amino acid side chains (ribbons and sticks).

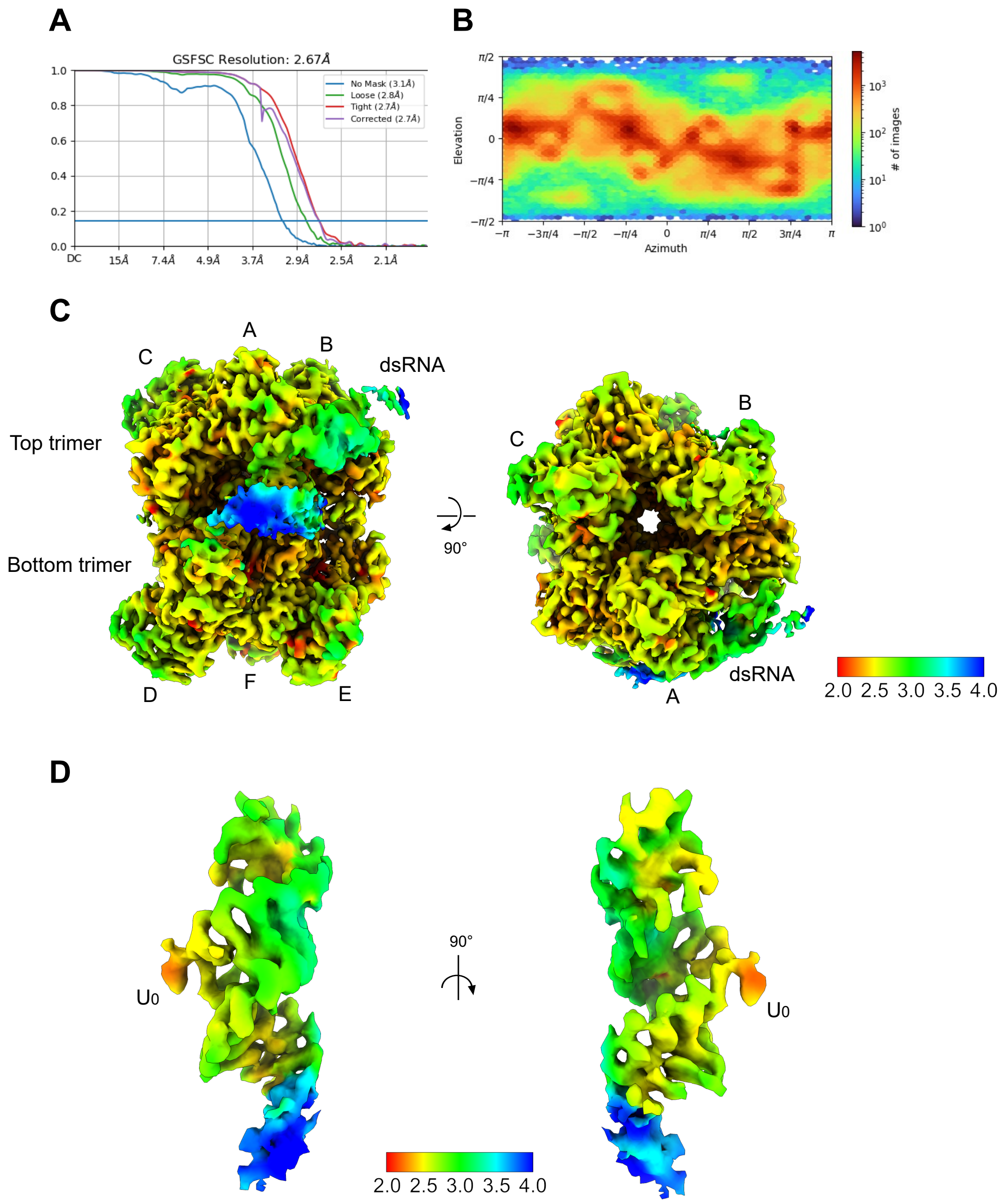


**Fig. S5.** Evaluation of the 3D reconstruction of Nsp15-RNA consensus structure. (A) Global resolution estimation of Nsp15-RNA. (B) Angular distribution plot of the particles of Nsp15-RNA. (C and D) Local resolution evaluation of the Nsp15-RNA (C) and its segmented RNA (D). Resolution estimation is based on the gold standard Fourier shell correlation (FSC) coefficient of 0.143 criteria.

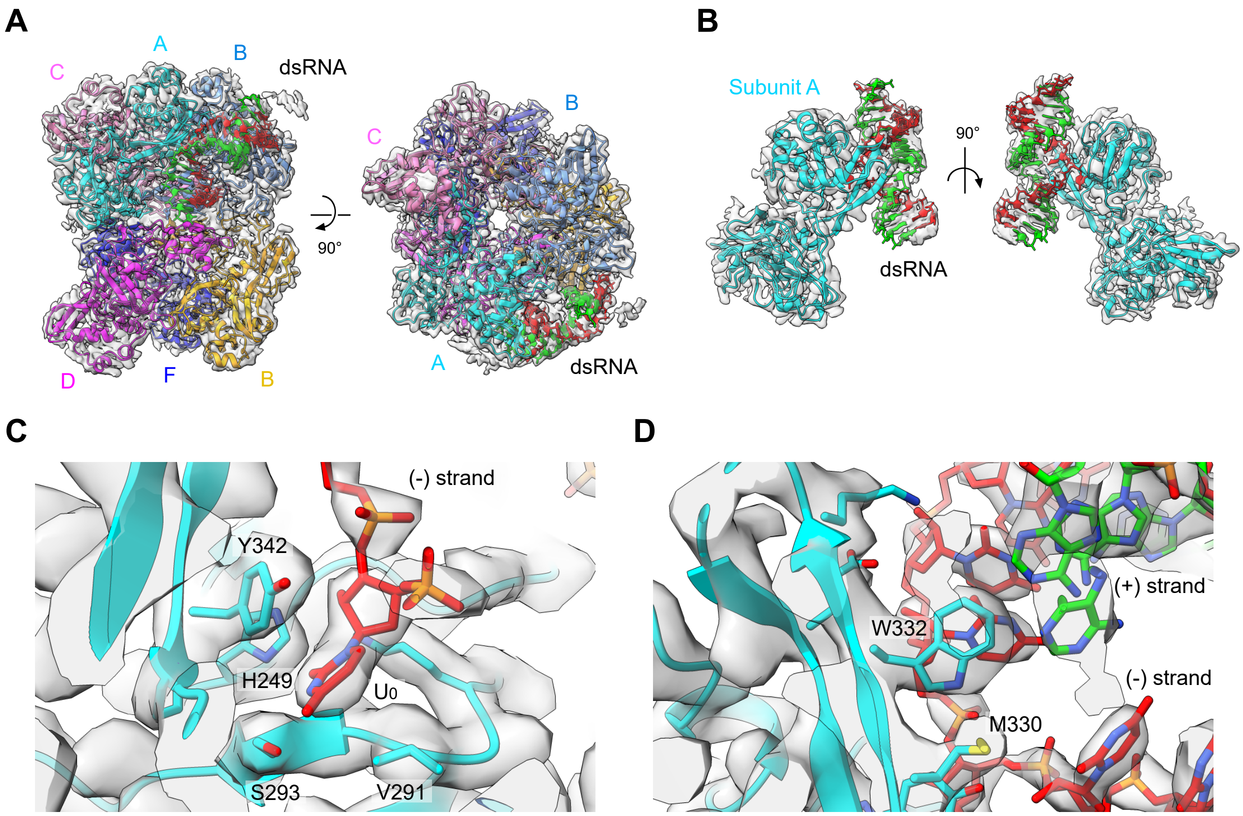


**Fig. S6.** Regions of the cryoEM density maps superposed with atomic models of the components of the Nsp15-RNA complex. Overall cryoEM density of Nap15-RNA (A), segmented cryoEM densities of the protomer A with the bound RNA (B), a local region around the $U_0$ base (C), and a local region around the base-flipping residues W332 and M330 (D) are shown as semi-transparent surfaces superposed with atomic models of amino acid side chains (ribbons and sticks).

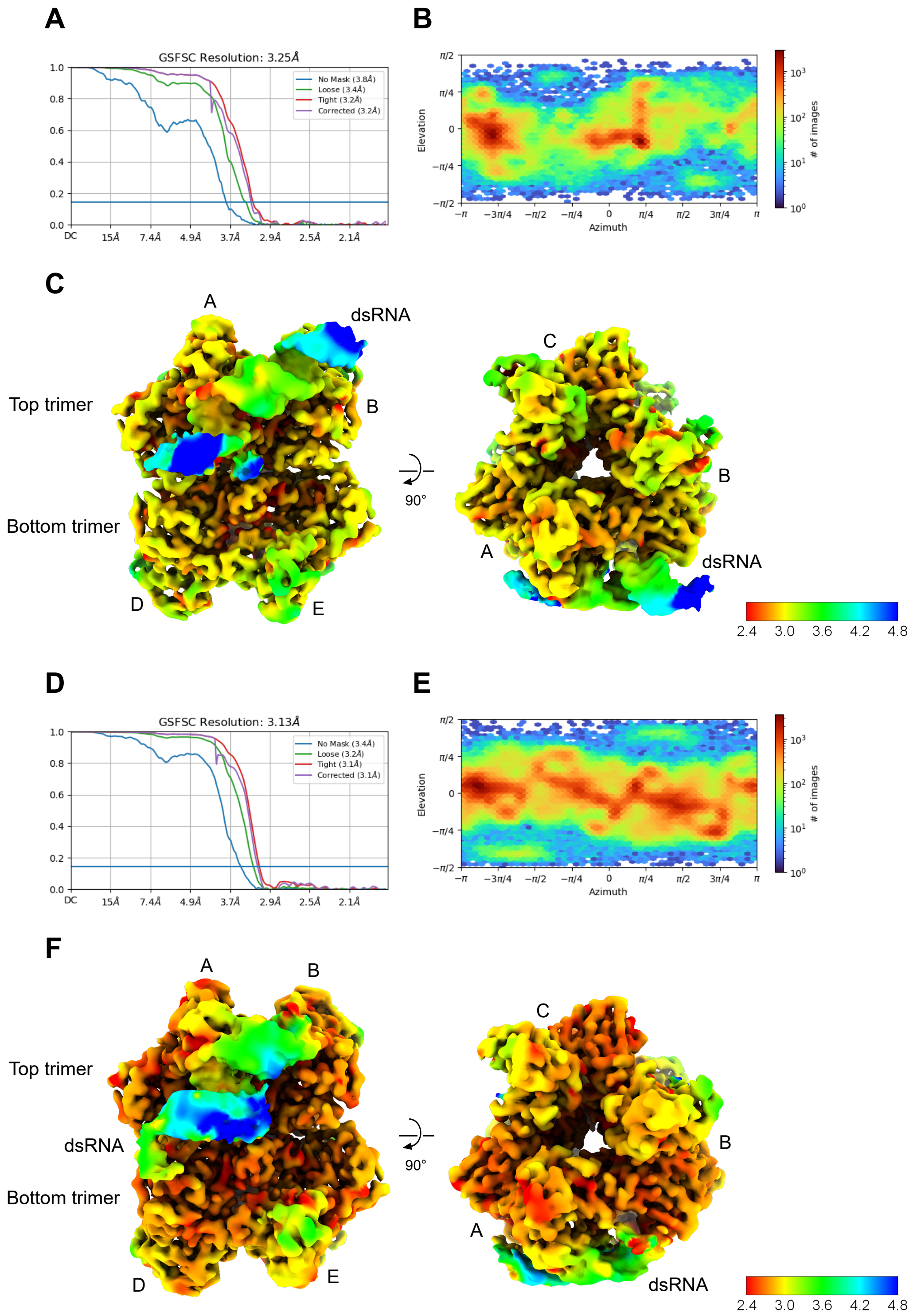


**Fig. S7.** Evaluation of the 3D reconstruction of two states of the Nsp15-RNA complex. (A and D) Global resolution estimation of Nsp15-RNA state 1 (A) and state 2 (D). (B and E) Angular distribution plot of the particles of Nsp15-RNA state 1 (B) and state 2 (E). (C and F) Local resolution evaluation of the Nsp15-RNA state 1 (C) and state 2 (F). Resolution estimation is based on the gold standard Fourier shell correlation (FSC) coefficient of 0.143 criteria.

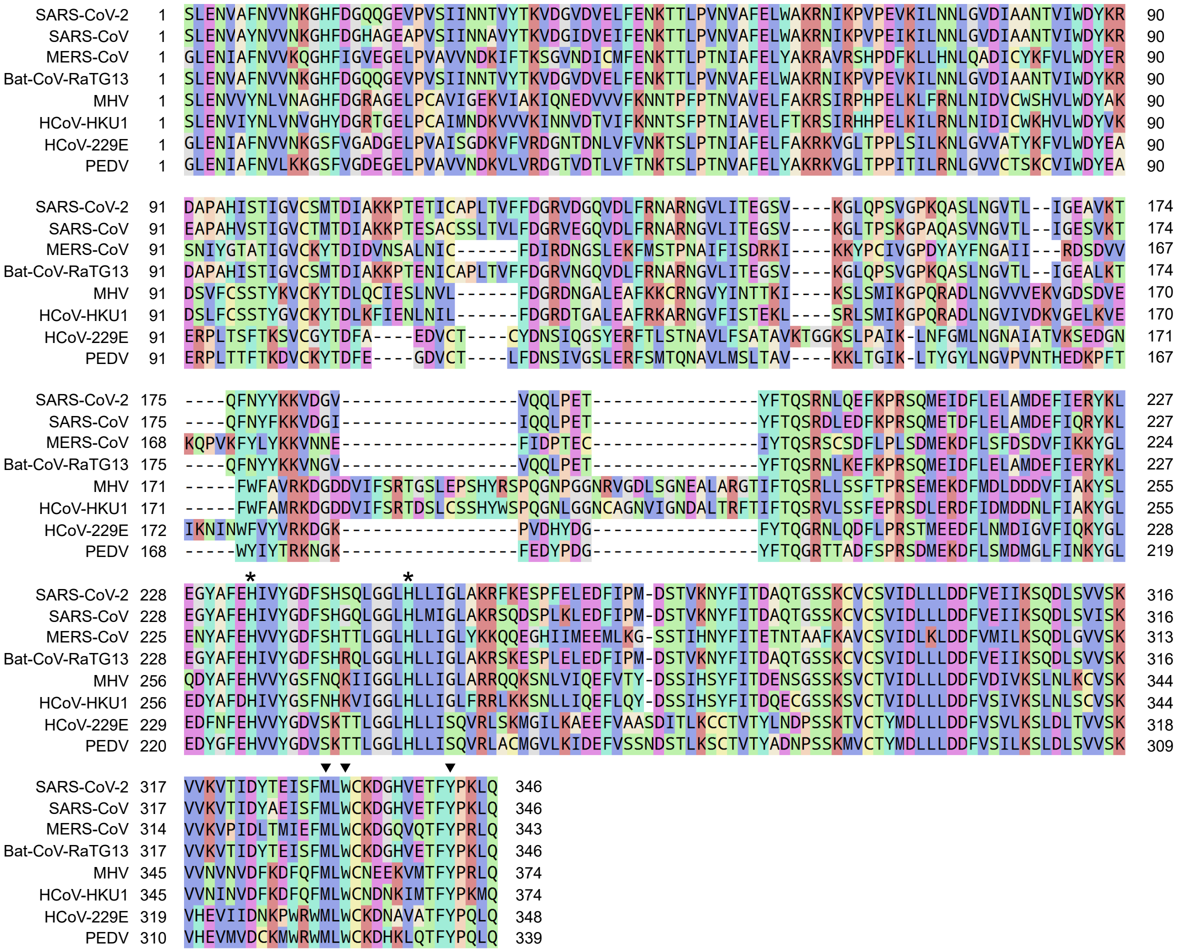


**Fig. S8. Sequence alignment of coronavirus Nsp15.** The functionally important residues in SARS-CoV-2 Nsp15 are marked for comparison with other coronavirus Nsp15s. Amino acid residues responsible for base-flipping are marked with triangles. Active site histidine residues are marked with asterisks. MERS-CoV, Middle East respiratory syndrome coronavirus; MHV, mouse hepatitis virus; PEDV, porcine epidemic diarrhea virus.

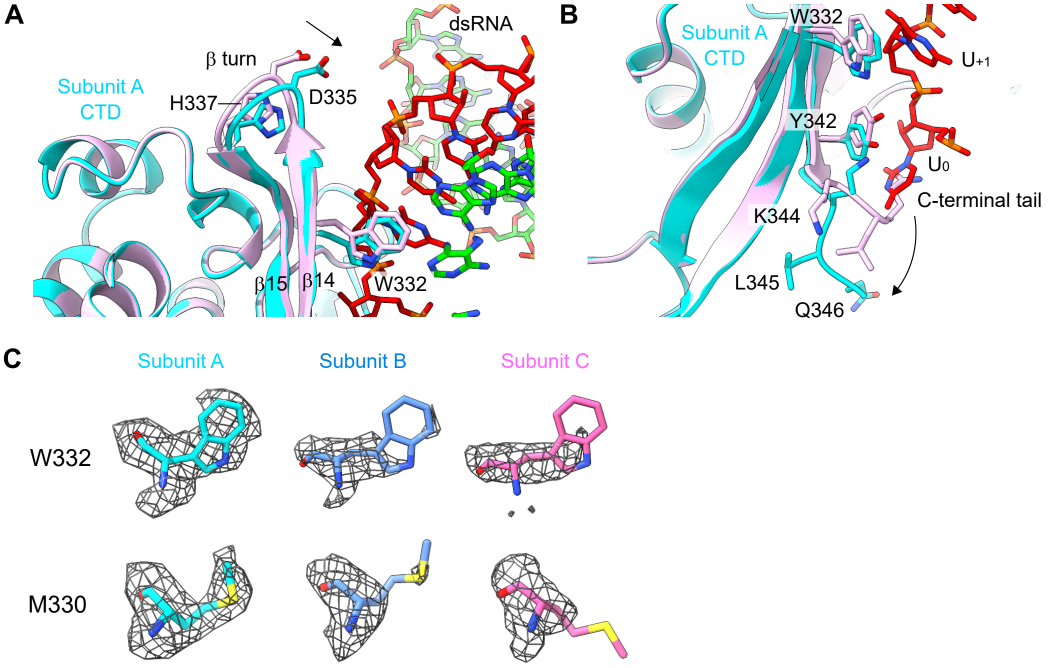


**Fig. S9.** Structural remodeling of Nsp15 upon RNA binding. (A) Fine-adjustment of the position of β14-tun-β15 region of subunit A's CTD. (B) Structural remodeling of the C-terminal tail of subunit A. C-terminal residues -$^{344}$KLQ$^{346}$ swings away from the active site center upon RNA binding. (C) The comparison of the density of W332 and M330 in different subunits at the same iso-surface threshold level (threshold: 0.32). The densities of subunit A show the clearer side-chain features of both W332 and M330 than those of the other subunits.
